# Genomic prediction of coronary heart disease

**DOI:** 10.1101/041483

**Authors:** Gad Abraham, Aki S. Havulinna, Oneil G. Bhalala, Sean G. Byars, Alysha M. De Livera, Laxman Yetukuri, Emmi Tikkanen, Markus Perola, Heribert Schunkert, Eric J. Sijbrands, Aarno Palotie, Nilesh J. Samani, Veikko Salomaa, Samuli Ripatti, Michael Inouye

**Affiliations:** Centre for Systems Genomics, School of BioSciences, The University of Melbourne, Parkville 3010, Victoria, Australia; Department of Pathology, The University of Melbourne, Parkville, Victoria 3010, Australia; National Institute for Health and Welfare, Helsinki, Finland; Centre for Epidemiology and Biostatistics, Melbourne School of Population and Global Health, The University of Melbourne, Parkville, Victoria 3010, Australia; Institute for Molecular Medicine Finland (FIMM), University of Helsinki, Helsinki, Finland; Deutsches Herzzentrum München, Klinik für Herz- und Kreislauferkrankungen, Munich, Germany; Department of Internal Medicine, Erasmus Medical Center, Rotterdam 3000 CA, The Netherlands; Analytic and Translational Genetics Unit, Department of Medicine, Massachusetts General Hospital, Boston, Massachusetts, USA; Program in Medical and Population Genetics, Broad Institute of Harvard and MIT, Cambridge, Massachusetts, USA; Psychiatric & Neurodevelopmental Genetics Unit, Department of Psychiatry, Massachusetts General Hospital, Boston, Massachusetts, USA; Department of Cardiovascular Sciences, University of Leicester, BHF Cardiovascular Research Centre, Glenfield Hospital, Groby Rd., Leicester, LE3 9QP, United Kingdom; National Institute for Health Research Leicester Cardiovascular Biomedical Research Unit, Glenfield Hospital, Groby Road, Leicester, LE3 9QP, United Kingdom; Wellcome Trust Sanger Institute, Wellcome Trust Genome Campus, Hinxton, Cambridge, United Kingdom; Department of Public Health, University of Helsinki, Helsinki, Finland

## Abstract

**Background:** Genetics plays an important role in coronary heart disease (CHD) but the clinical utility of a genomic risk score (GRS) relative to clinical risk scores, such as the Framingham Risk Score (FRS), is unclear.

**Methods:** We generated a GRS of 49,310 SNPs based on a CARDIoGRAMplusC4D Consortium meta-analysis of CHD, then independently tested this using five prospective population cohorts (three FINRISK cohorts, combined n=12,676, 757 incident CHD events; two Framingham Heart Study cohorts (FHS), combined n=3,406, 587 incident CHD events).

**Results:** The GRS was strongly associated with time to CHD event (FINRISK HR=1.74, 95% CI 1.61-1.86 per S.D. of GRS; Framingham HR=1.28, 95% CI 1.18-1.38), and was largely unchanged by adjustment for clinical risk scores or individual risk factors, including family history. Integration of the GRS with clinical risk scores (FRS and ACC/AHA13 score) improved prediction of CHD events within 10 years (meta-analysis C-index: +1.5-1.6%, *P*<0.001), particularly for individuals ≥60 years old (meta-analysis C-index: +4.6-5.1%, *P*<0.001). Men in the top 20% of the GRS had 3-fold higher risk of CHD by age 75 in FINRISK and 2-fold in FHS, and attaining 10% cumulative CHD risk 18y earlier in FINRISK and 12y earlier in FHS than those in the bottom 20%. Furthermore, high genomic risk was partially compensated for by low systolic blood pressure, low cholesterol level, and non-smoking.

**Conclusions:** A GRS based on a large number of SNPs substantially improves CHD risk prediction and encodes decades of variation in CHD risk not captured by traditional clinical risk scores.

## Introduction

Early and accurate identification of individuals with increased risk of coronary heart disease (CHD) is critical for effective implementation of preventative lifestyle modifications and medical interventions, such as statin treatment^1,2^. To this end, risk scores such as the Framingham Risk Score (FRS)^3^ and the American College of Cardiology / American Heart Association 2013 risk score (ACC/AHA13)^1^, based on clinical factors and lipid measurements, have been developed and are widely used. Although the scores can identify individuals at very high risk, a large proportion of individuals developing CHD during the next 10 years remain unidentified. In particular, they do not provide sufficient discrimination at a younger age when implementation of preventative measures is likely to provide the greatest long-term benefit.

Genetic factors have long been recognized to make a substantial contribution to CHD risk^4^. Although a positive family history is an independent risk factor for CHD, it may not completely and solely capture genetic risk. Recently, genome-wide association studies (GWAS) have identified 56 genetic loci associated with CHD at genome-wide significance^5-9^. Studies of the predictive power of the top single nucleotide polymorphisms (SNPs) at some of these loci either individually or in combination have typically shown small improvements in CHD risk prediction^10-19^, probably because together these variants only explain less than 20% of CHD heritability^8^. As demonstrated recently for other traits such as height and BMI^20,21^, the majority of unexplained heritability is likely hidden amongst the thousands of SNPs that did not reach genome-wide significance. Indeed, recent advances have shown that genomic prediction models that consider all available genetic variants can efficiently stratify those at increased risk of complex disease^22,23^. To leverage the maximum amount of information, we examined whether a genomic risk score (GRS) comprising a large number of SNPs, including those with less than genome-wide significance, could produce clinically relevant predictive power for CHD risk.

## Methods

A summary of the key methods for the study is given here. Additional details are provided in the **Supplementary Appendix**.

### Study design

This study consisted of two main stages (**Supplementary Figure S1**). In stage 1, we utilized the large-scale CHD genetic association dataset assembled by the CARDIoGRAMplusC4D consortium^8^ (downloaded from http://www.cardiogramplusc4d.org) to construct a GRS. Briefly, CARDIoGRAMplusC4D tested associations of 79,128 SNPs in 63,746 CHD cases and 130,681 controls, including 6,222 SNPs that had shown nominal association with CHD in a prior GWAS meta-analysis^24^. We extracted information on all these SNPs together with their associated CARDIoGRAMplusC4D weights (log odds) and used two case/control datasets, the Wellcome Trust Case/Control Consortium Coronary Artery Disease dataset (WTCCC-CAD)^5^ (1,926 cases and 2,938 controls) and the MIGen case/control dataset^7^ (Harps subset) (531 cases and 488 controls), to optimize the predictive value of this set of SNPs using linkage-disequilibrium (LD) thinning to derive a GRS. In stage 2, we assessed the predictive accuracy of this GRS with incident CHD events in the FINRISK and the Framingham Heart Study cohorts, comparing it to traditional risk factors and clinical risk scores. Secondary validation was also performed in the ‘Association of CHD Risk in a Genome-wide Old-versus-young Setting’ (ARGOS) study^29^ familial hypercholesterolemia study. Details of the final GRS used are available at http://www.inouyelab.org.

### Prospective Study cohorts

We utilized two sets of prospective cohorts: (i) FINRISK, consisting of three prospective cohorts from Finland with 10-20 years of follow-up, from collections 1992, 1997, and 2002 (FR92, FR97, and FR02, respectively)^25^, and (ii) the Framingham Heart Study (FHS)^26-28^, with individuals of Western and Southern European ancestry taken from the Original and Offspring cohorts with 40-48 years of follow-up. In total, the FINRISK consisted of n=12,676 individuals and the FHS of n=3,406 individuals, all of whom had the requisite data and were independent of the CARDIoGRAMplusC4D stage-2 meta-analysis utilized to generate the GRS. Details of the cohorts are given in Table 1. The cohorts have been genome-wide SNP genotyped (Illumina BeadArrays for FINRISK and Affymetrix 500K for FHS) and further imputed to the 1000 Genomes reference panel (**Supplementary Methods**). After genotype imputation and quality control, 69,044 autosomal SNPs of the 79,128 CARDIoGRAMplusC4D SNPs were available for subsequent analyses in the FINRISK, and 78,058 autosomal SNPs available in FHS.

**Table 1:**
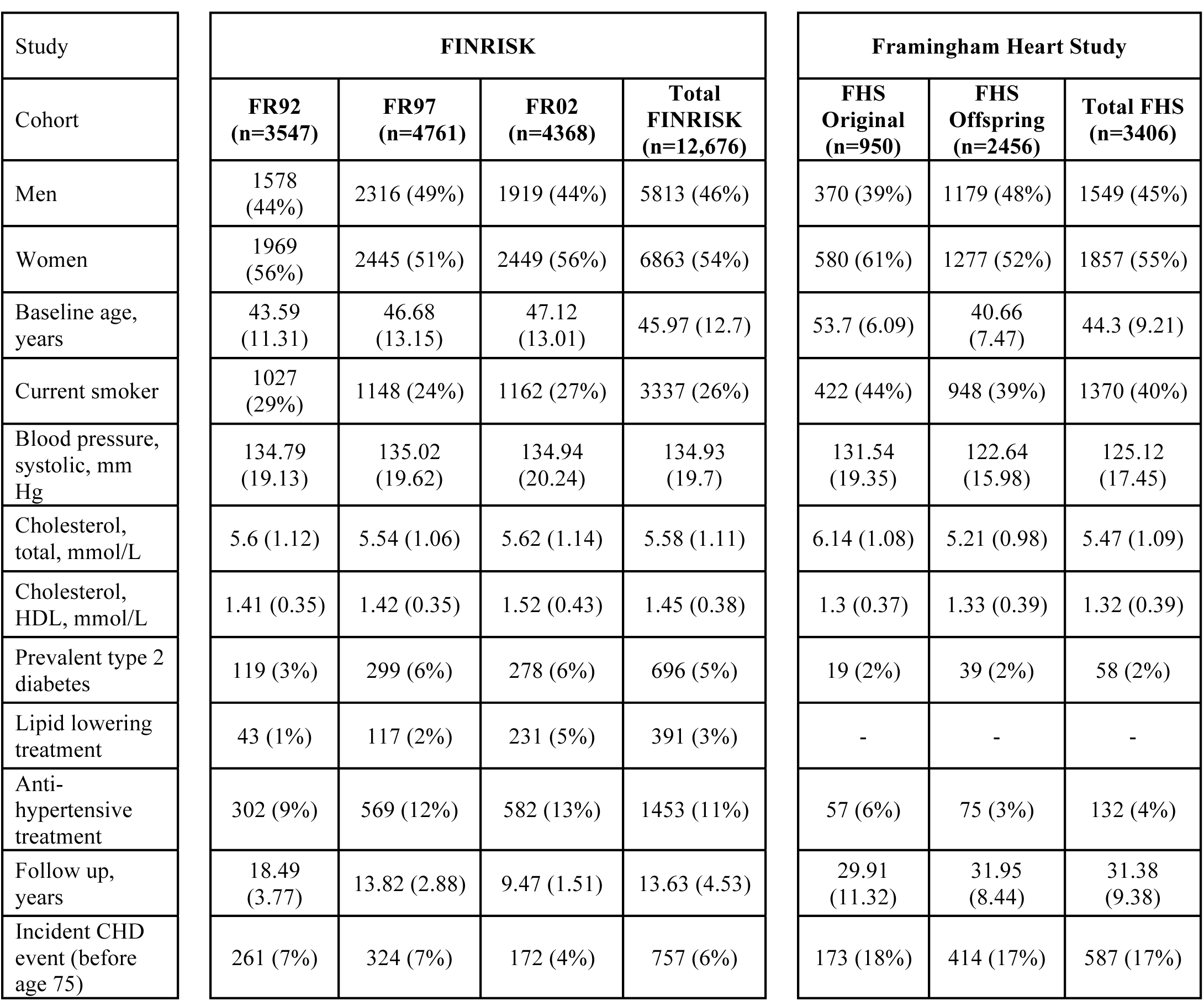
Characteristics of the FINRISK and FHS cohorts. Categorical variables are shown as counts and percentages, continuous variables (age, follow-up time, cholesterol, and blood pressure) as means and standard deviations. Sample sizes are for participants with GWAS data after quality control and all other exclusions. Lipid lowering treatments were not assessed in FHS due to an insufficient number of exams with this information.

The outcome of interest in FINRISK was primary incident coronary heart disease (CHD) event, defined as myocardial infarction (MI), a coronary revascularization procedure, or death from CHD, before age 75y (**Supplementary Methods**). Individuals with prevalent cardiovascular disease (CVD) at baseline were excluded from the analysis. We censored events for individuals with an attained age of >75y, as not all FINRISK cohorts had sufficient numbers of CHD events beyond that age. In FHS, we used the FHS definition of CHD, which included recognized/unrecognized MI, death from CHD, angina pectoris, and coronary insufficiency (**Supplementary Methods**). For FHS analyses, we excluded the Third cohort due to insufficient follow-up time (median 6.6 years, max 9.5 years), and for each individual we selected the first exam yielding the most complete set of non-missing clinical risk factors (for Original cohort: exams 9, 10, 15, 20, and 22; for Offspring cohort: exams 1, 3, 4, 5, and 6). FHS individuals with prevalent CHD or <30 years of age at baseline were excluded, and for consistency with the FINRISK analysis, a censoring age of 75y was also applied to the FHS analyses.

Secondary validation of the GRS was performed in the ARGOS study, a Dutch case/control dataset where all individuals had familial hypercholesterolemia (248 young cases with early CHD, 216 elderly controls without CHD), imputed to 1000 Genomes reference panel (74,135 SNPs of the 79,128 CARDIoGRAMplusC4D SNPs were available; **Supplementary Methods**).

### Statistical analysis

Clinical risk scores were based on the FRS (Adult Treatment Panel III) for CHD and/or MI^3^, and the recent ACC/AHA13 risk score^1^ for atherosclerotic cardiovascular disease (ASCVD; see **Supplementary Tables S1** and **S2** for components and derivation of the scores).

GRSs were generated via thinning the CARDIoGRAMplusC4D SNPs by linkage disequilibrium (LD) thresholds and evaluated using logistic regression and area under receiver-operating characteristic curve (AUC) for each threshold (**Supplementary Figure S2**). To avoid overfitting we only used weights (log odds) from the CARDIoGRAMplusC4D stage-2 meta-analysis, which were not based on the WTCCC-CAD or MIGen studies (see **Supplementary Methods**). We combined the estimates for WTCCC and MIGen-Harps using fixed-effects inverse-variance weighted meta-analysis. We compared the performance of our GRS with those of Tikkanen et al.^11^ and of Mega et al.^30^, which were based on 28 and 27 SNPs associated with CHD at genome-wide significance, respectively.

For analysis of FINRISK, we used Cox proportional hazard models to evaluate the association of the GRS with time to incident CHD events, stratifying by sex and adjusting for geographic location and cohort, using age as the time scale. Secondary analyses adjusted for one of the clinical risk scores (FRS or ACC/AHA13), or individual baseline variables and known risk factors (cohort, geographical location, prevalent type-2 diabetes, log total cholesterol, log HDL, log systolic BP, smoking status, lipid treatment, and family history). Family history in FINRISK was self-reported and was defined as having a 1^st^-degree relative who had experienced myocardial infarction before age 60. For FHS, we evaluated the association of the GRS with incident CHD using Cox proportional hazard models, stratifying by sex and adjusting for cohort (Original or Offspring), using age as the time scale. Family history was not available for both FHS cohorts and thus not considered in FHS analyses. Survival analyses allowing for competing risks were performed using the Aalen-Johansen estimator of survival and cause-specific Cox models (**Supplementary Methods).** Model discrimination of incident CHD event was evaluated in three groups of individuals: (i) all individuals (n=12,676 in FINRISK, n=3,406 in FHS), (ii) individuals aged <60 years at baseline (n=10,606 in FINRISK, n=3,218 in FHS), and (iii) individuals aged ≥60 years at baseline (n=2,070 in FINRISK, n=188 in FHS).

Discrimination of incident CHD events within 10 years was assessed using Harrell’s C-index, and the difference in C-index between two models was assessed using the correlated jackknife test. Competing risk analyses were performed using the Aalen-Johansen empirical estimator of cumulative incidence and cause-specific Cox proportional hazard models. Risk reclassification was evaluated using continuous Net Reclassification Improvement, categorical Net Reclassification Improvement, and Integrated Discrimination Improvement. Meta-analysis of the discrimination statistics was performed using fixed-effect inverse-variance weighting. Additional details on the statistical methods are provided in the **Supplementary Methods**.

### Role of funding source

The sponsors had no role in the conduct or interpretation of the study. The corresponding authors had full access to all data in the study and had final responsibility for the decision to submit for publication.

## Results

To construct an optimized GRS using the WTCCC and MIGen-Harps datasets, we first generated a series of GRSs, starting with the 79,128 CARDIoGRAMplusC4D SNPs then progressively lowering the *r*^2^ threshold for LD to reduce the redundancy of predictive information and corresponding number of SNPs in the score (**Methods** and **Supplementary Figure S1**). An *r^2^* threshold of 0.7 provided optimal discrimination of CHD cases and controls (WTCCC and MIGen-Harps meta-analysis OR=1.70 per S.D. of GRS, 95% CI 1.61-1.80; meta-analysis AUC=0.64, 95% CI 0.63-0.66), corresponding to 49,310 SNPs in WTCCC (**Supplementary Figure S2**). Of these 49,310 SNPs, 85.9% (42,364 SNPs) and 95% (46,773 SNPs) were available in the FINRISK and FHS, respectively.

In analyses of incident CHD as a binary outcome in FINRISK, the 49K GRS showed similar odds ratios to WTCCC and MIGen-Harps (OR=1.72, 95% CI 1.59-1.86, per S.D.), but led to higher discrimination than in WTCCC and MIGen-Harps (AUC=0.73, 95% CI 0.71-0.75). In FHS, the association was weaker, OR=1.30 (95% CI 1.19-1.43, per S.D.), with AUC=0.65 (95% CI 0.62-0.67).

Using survival analyses of time to incident CHD, within FINRISK the GRS had stronger association with CHD (HR=1.74, 95% CI 1.61-1.86, per S.D.) than the 28 SNP score studied by Tikkanen et al.^11^ (HR=1.21, 95% CI 1.13-1.30, per S.D.) or the 27 SNP score used by Mega et al.^30^ (HR=1.21, 95% CI 1.12-1.30 per S.D.) (**Supplementary Results**). In FHS, the GRS showed weaker but statistically significant association with CHD (HR=1.28 per S.D. of the GRS, 95% CI 1.18-1.38). The fixed-effect meta-analysis estimate for the GRS combining FINRISK and FHS was HR=1.66 (95% CI 1.55-1.78), however, heterogeneity was high (*I*^2^=89.2%, *P*=0.0023). For both FINRISK and FHS, the GRS showed improved prediction for incident CHD over the other risk scores composed of smaller numbers of SNPs (**Supplementary Results** and **Supplementary Table 3**).

In both FINRISK and FHS, the hazard ratios for GRS were not substantially attenuated by adjusting for FRS or ACC/AHA13 clinical risk scores, lipid treatment at baseline, other established risk factors (including family history in FINRISK), or 5 principal components of the genotypes (**Supplementary Figures S3** and **S4**). The correlation between GRS and either FRS or ACC/AHA13 scores was close to zero with almost none of the variation in GRS explained by either clinical risk score (in both FINRISK and FHS, *r*^2^<0.004 between GRS and either FRS and ACC/AHA13; **Supplementary Figure S5**). To further test that the CHD risk conferred by the GRS was largely independent of the effects of cholesterol, we further validated the GRS in the ARGOS familial hypercholesterolemia study, with comparable results to those obtained in WTCCC/MIGen (OR=1.49, 95% CI 1.21-1.84 per S.D. of the GRS, adjusted for sex and 5 principal components) (**Supplementary Methods**).

To assess the potential clinical utility of the GRS, we compared its performance in discrimination of time to CHD event (C-index) with that of family history, the previously published 28-SNP Tikkanen score^11^ the 27-SNP Mega score^30^, and the FRS and ACC/AHA13 clinical risk scores. We also assessed the incremental value of the GRS on top of the clinical risk scores. In both FINRISK and FHS, addition of GRS to either FRS or ACC/AHA13 scores provided statistically significant improvements in C-index, in FINRISK: +1.7% (*P*<10^−6^) and +1.6% (*P*<10^−6^) for FRS and ACC/AHA13, respectively; in FHS: +1.1% (*P*<0.0443) and +1.1% (*P*<0.0344) for FRS and ACC/AHA13, respectively (Figure 1). Overall, fixed-effects meta-analysis of the two studies showed that GRS improved the C-index by +1.6% (95% CI 0.01-0.02, *P*<10^−6^; heterogeneity: *I*^2^=2.2%, *Q*=1.02, *P*=0.312) for FRS and GRS combined (FRS+GRS) over FRS alone and, similarly, +1.5% (95% CI 0.009-0.02, *P*<10^−6^; heterogeneity: *I*^2^=0%, *Q*=0.78, *P*=0.378) for ACC/AHA13+GRS over ACC/AHA13 alone (Figure 1). Larger increases in C-index were observed among older individuals, with the C-index of FRS+GRS compared to FRS alone increasing by 5.1% in individuals aged ≥60 years at baseline, while individuals aged <60 years at baseline showed C-index gains of 1.4% (**Supplementary Figure S6**). Within FINRISK, the GRS had higher C-index than family history (+1.9%, *P*<1.3×10^−6^). In meta-analysis, the GRS had higher C-index than the published scores of Tikkanen et al. and Mega et al. (meta-analysis C-index improvement +1.8-2.1%, both *P*<1×10^−6^) (**Supplementary Results** and **Supplementary Table 3**).

**Figure 1:**
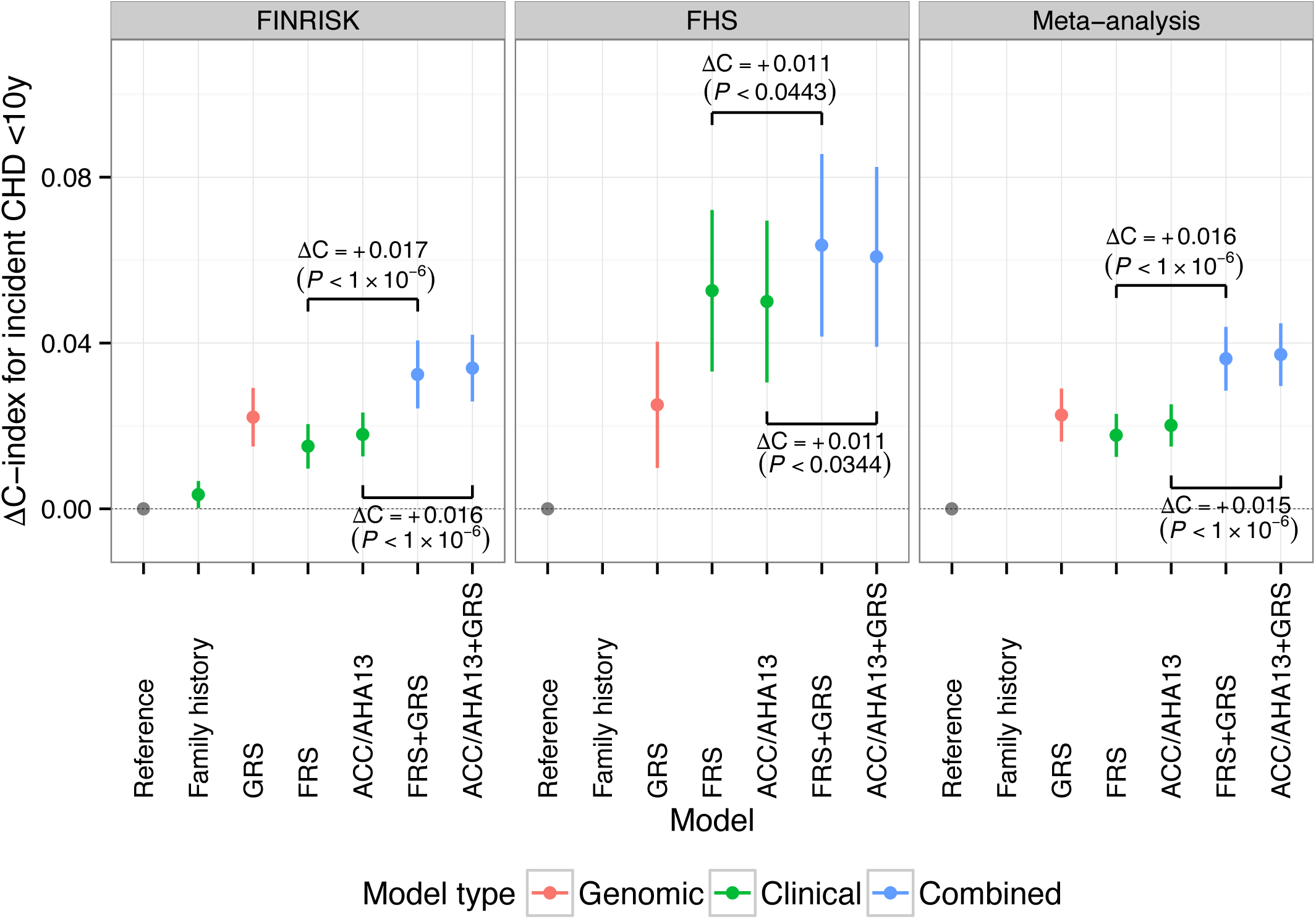
Difference in C-index (95% CI) for time to incident CHD event within 10y, relative to the reference model in the FINRISK and FHS cohorts. Reference models used age as the time scale, stratified by sex (FINRISK: adjusted for cohort and geographic location; FHS: adjusted for cohort). Family history was not available for all of the FHS cohorts and thus not considered here. P-values are from the correlated jackknife test.

We next assessed if the GRS improved the individual risk reclassification when added to clinical risk scores. Analyses within FINRISK and FHS are given in Table 2 for FRS and in **Supplementary Table S4** for ACC/AHA13. Overall, meta-analysis of the two datasets showed that the categorical Net Reclassification Improvement (NRI) was 0.1 for both the FRS+GRS and ACC/AHA13+GRS, respectively (*P*<0.0001; **Supplementary Figure S7**). Meta-analysis of continuous NRI was 0.344 (*P*<0.001) and 0.334 (*P*<0.001) for the FRS+GRS and ACC/AHA13+GRS, respectively **(Supplementary Figure S8**). Meta-analysis of IDI scores showed gains of 0.01 (*P*<0.001) and 0.009 (*P*<0.001) for FRS+GRS and ACC/AHA13+GRS, respectively, however IDI scores showed high heterogeneity across FINRISK and FHS (*I*^2^>97%, *P*<0.0001, **Supplementary Figure S9**).

**Table 2:**
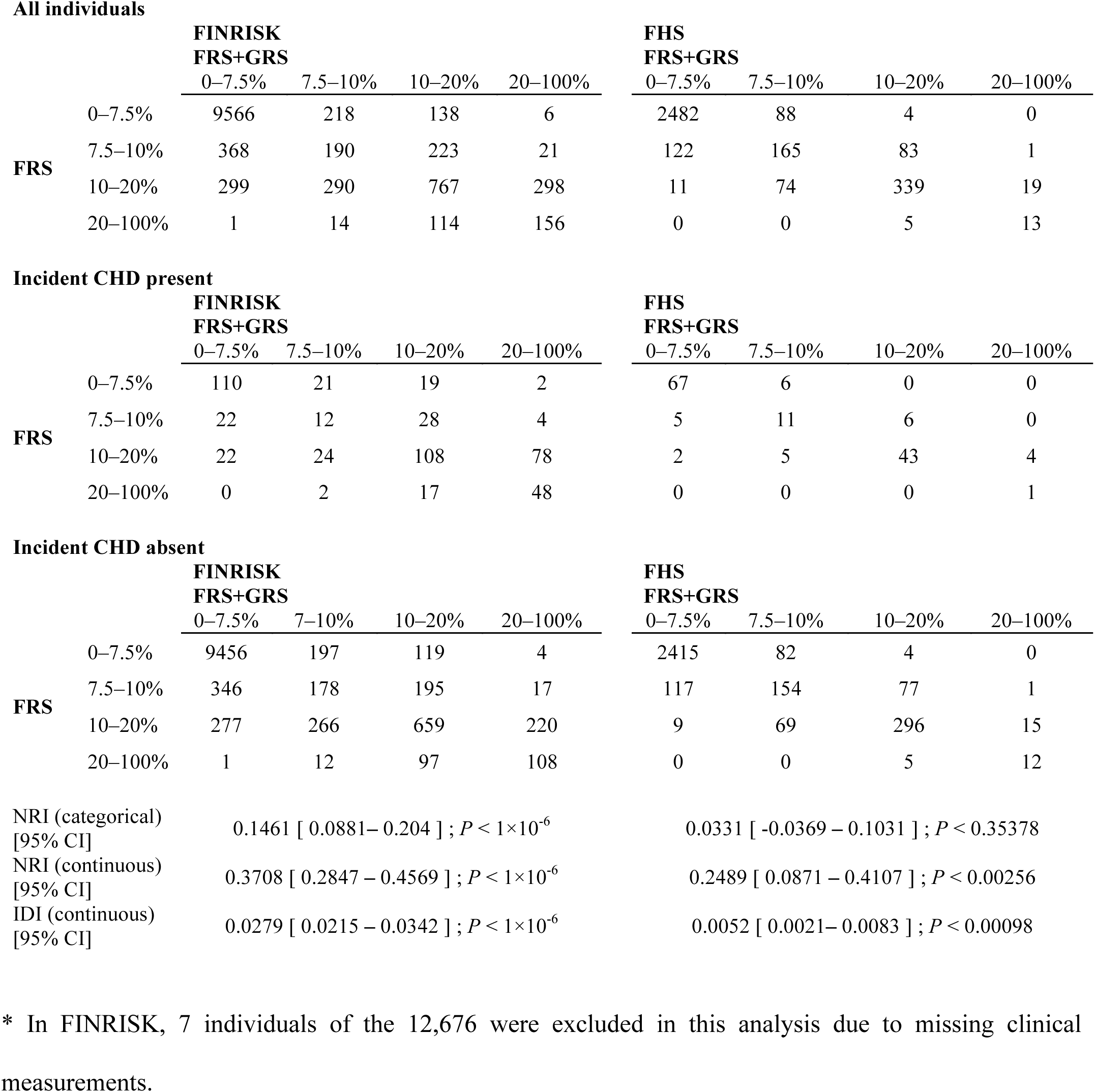
Reclassification of incident CHD event risk within 10 years for combined FRS + GRS compared to FRS only, in the FINRISK and FHS cohorts.

**All individuals**
|  |  | **FINRISK**<br>**FRS+GRS** |  |  |  | **FHS**<br>**FRS+GRS** |  |  |  |
| --- | --- | --- | --- | --- | --- | --- | --- | --- | --- |
|  |  | 0–7.5% | 7.5–10% | 10–20% | 20–100% | 0–7.5% | 7.5–10% | 10–20% | 20–100% |
| **FRS** | 0–7.5% | 9566 | 218 | 138 | 6 | 2482 | 88 | 4 | 0 |
|  | 7.5–10% | 368 | 190 | 223 | 21 | 122 | 165 | 83 | 1 |
|  | 10–20% | 299 | 290 | 767 | 298 | 11 | 74 | 339 | 19 |
|  | 20–100% | 1 | 14 | 114 | 156 | 0 | 0 | 5 | 13 |

**Incident CHD present**
|  |  | **FINRISK**<br>**FRS+GRS** |  |  |  | **FHS**<br>**FRS+GRS** |  |  |  |
| --- | --- | --- | --- | --- | --- | --- | --- | --- | --- |
|  |  | 0–7.5% | 7.5–10% | 10–20% | 20–100% | 0–7.5% | 7.5–10% | 10–20% | 20–100% |
| **FRS** | 0–7.5% | 110 | 21 | 19 | 2 | 67 | 6 | 0 | 0 |
|  | 7.5–10% | 22 | 12 | 28 | 4 | 5 | 11 | 6 | 0 |
|  | 10–20% | 22 | 24 | 108 | 78 | 2 | 5 | 43 | 4 |
|  | 20–100% | 0 | 2 | 17 | 48 | 0 | 0 | 0 | 1 |

**Incident CHD absent**
|  |  | **FINRISK**<br>**FRS+GRS** |  |  |  | **FHS**<br>**FRS+GRS** |  |  |  |
| --- | --- | --- | --- | --- | --- | --- | --- | --- | --- |
|  |  | 0–7.5% | 7–10% | 10–20% | 20–100% | 0–7.5% | 7.5–10% | 10–20% | 20–100% |
| **FRS** | 0–7.5% | 9456 | 197 | 119 | 4 | 2415 | 82 | 4 | 0 |
|  | 7.5–10% | 346 | 178 | 195 | 17 | 117 | 154 | 77 | 1 |
|  | 10–20% | 277 | 266 | 659 | 220 | 9 | 69 | 296 | 15 |
|  | 20–100% | 1 | 12 | 97 | 108 | 0 | 0 | 5 | 12 |

|  |  |  |
| --- | --- | --- |
| NRI (categorical)<br>[95% CI] | 0.1461 [ 0.0881 – 0.204 ] ; $P < 1 \times 10^{-6}$ | 0.0331 [ -0.0369 – 0.1031 ] ; $P < 0.35378$ |
| NRI (continuous)<br>[95% CI] | 0.3708 [ 0.2847 – 0.4569 ] ; $P < 1 \times 10^{-6}$ | 0.2489 [ 0.0871 – 0.4107 ] ; $P < 0.00256$ |
| IDI (continuous)<br>[95% CI] | 0.0279 [ 0.0215 – 0.0342 ] ; $P < 1 \times 10^{-6}$ | 0.0052 [ 0.0021 – 0.0083 ] ; $P < 0.00098$ |

\* In FINRISK, 7 individuals of the 12,676 were excluded in this analysis due to missing clinical measurements.

The FRS+GRS model also showed consistently higher positive predictive value (PPV) at any given negative predictive value (NPV) for 10-year incident CHD events (**Supplementary Figure S10**). For example, in FINRISK, at an NPV of 96% the FRS+GRS achieved a PPV of 40%, compared with a PPV of 23% for FRS alone. In FHS, PPV for FRS+GRS and ACC/AHA13+GRS models was 12-13% at the same NPV, compared with PPV of 10% for the FRS or ACC/AHA13 scores (**Supplementary Figure S10**).

We next examined how variation in genomic risk translated into differences in cumulative lifetime risk of CHD, using Kaplan-Meier estimates stratified by GRS quintiles for men and women separately (Figure 2). As expected, cumulative risk increased with age for both sexes, with men displaying higher absolute risk than women. In both sexes there were substantial differences in cumulative risk between GRS groups with 1.7-fold (in FHS) to 3.2-fold (in FINRISK) higher cumulative risk by age 75 in those in the top quintile of GRS versus bottom quintile. When considering clinically relevant levels of risk, FINRISK men in the top quintile of genomic risk achieved 10% cumulative risk 18 years earlier than those in the bottom quintile (ages 52 and 70, respectively), with a comparable difference of 12 years in FHS (ages 51 and 64). Women in the top quintile of genomic risk achieved 10% cumulative risk by age 69 (FINRISK) and 64 (FHS), whereas women in the bottom quintile did not achieve 10% risk by age 75 in FINRISK, or by age 73 in FHS. Estimated lifetime CHD risk in FINRISK showed no evidence of being affected by competing risks (incident CHD versus non-CHD death) (**Supplementary Methods** and **Supplementary Figure S11**). Similarly, a cause-specific competing-risk Cox analysis of the GRS in FINRISK, adjusting for geographical location and cohort, resulted in a similar hazard ratio as standard Cox analysis (HR=1.70, 95% CI 1.61-1.86).

**Figure 2:**
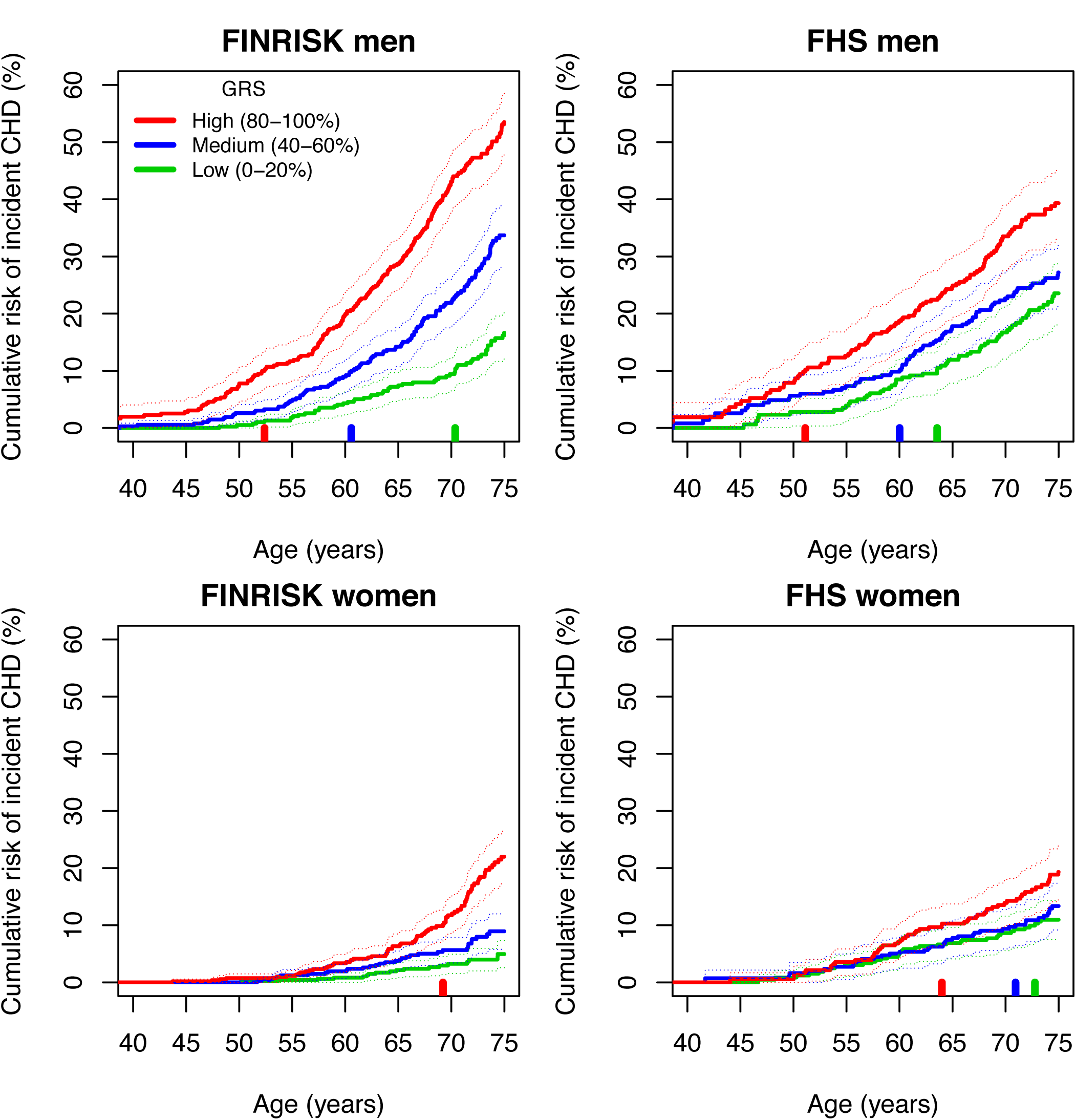
Kaplan-Meier cumulative risk of incident CHD event by genomic risk group for men and women in the FINRISK and FHS cohorts. Showing the cumulative risk in quintiles 0-20%, 40-60%, 80-100%. The vertical bars along the x-axis indicate the age at which each risk group attains a cumulative CHD risk of 10%. Dashed lines indicate 95% CI.

We next sought to investigate to what degree high genomic risk for CHD could be compensated for by low levels of clinical risk factors at baseline, and vice-versa. When considering baseline smoking status in both FINRISK and FHS, Kaplan-Meier analysis showed a substantial increase in cumulative risk of CHD in men who smoked and also in the top quintile of genomic risk, relative to either non-smokers or smokers at low genomic risk (Figure 3 for FINRISK and **Supplementary Figure S12** for FHS). Similar but weaker trends were observed for women in the top versus bottom quintiles of genomic risk. To test whether there was evidence for smoking affecting CHD hazard differently based on an individual’s genomic background, we used a Cox model allowing for an interaction term between the GRS and smoking; the interaction was not statistically significant in FINRISK (*P*=0.91) and FHS (*P*=0.49).

**Figure 3:**
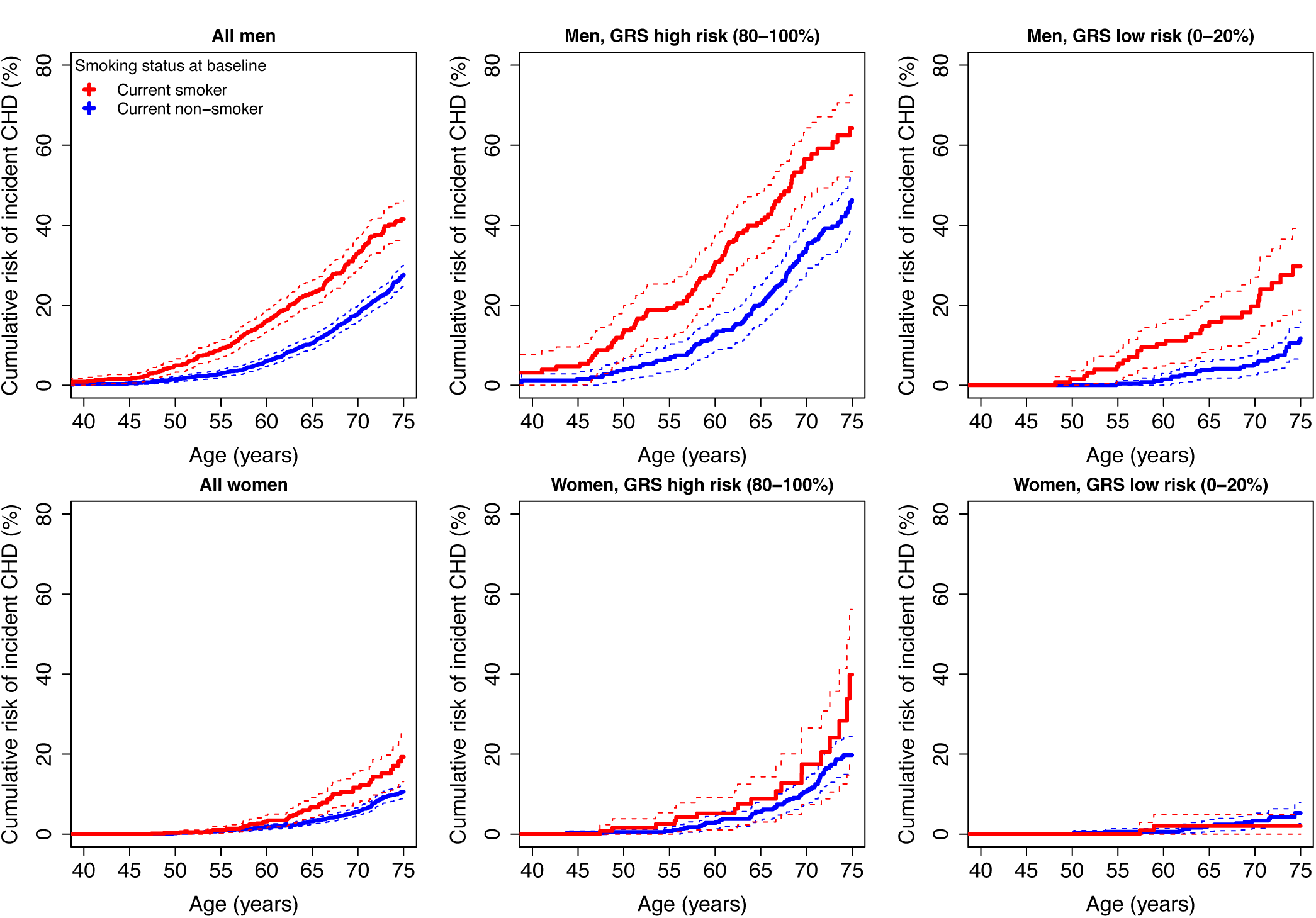
Kaplan-Meier curves for incident CHD event risk stratified by GRS quintiles and smoking status at baseline, for men and women in the FINRISK cohorts.

We also examined the potential compensatory effects of baseline systolic blood pressure and total cholesterol, divided as tertiles of high, medium, and low levels (**Supplementary Figures S13** and **S14**). For both systolic blood pressure and total cholesterol, we observed the expected trends in CHD risk for high, medium and low levels. However, males with high versus low levels of systolic blood pressure or total cholesterol showed greater absolute CHD risk if they were in the top versus bottom quintiles of genomic risk.

Notably, in both FINRISK and FHS, women in the bottom quintile of genomic risk showed smaller differences in cumulative CHD risk when stratified by smoking. For tertiles of systolic blood pressure or total cholesterol, low genomic risk women in FINRISK showed similarly small differences in risk, but the effects in FHS for this subgroup were not consistent. Cox models allowing for interactions between the GRS and systolic blood pressure or total cholesterol did not show statistically significant interactions in either FINRISK or FHS (*P*>0.2 for all).

## Discussion

We have generated a GRS for CHD based on 49,310 SNPs and, using three prospective FINRISK and two FHS prospective cohorts, demonstrated that the GRS is associated with incident CHD events independently of established and widely-used clinical risk scores or individual CHD risk factors, including family history. Secondary validation in a familial hypercholesterolemia study (ARGOS) showed that GRS was also associated with CHD in this group of high-risk individuals. Subsequently, combining the GRS with established risk scores improved 10-year CHD risk prediction in FINRISK and FHS. We have also shown that the GRS can be leveraged to achieve meaningful lifetime CHD risk stratification, and that the impact of traditional CHD risk factors such as smoking, blood pressure, and cholesterol, vary substantially depending on the underlying genetic risk, thus offering the potential for both earlier and more targeted preventative efforts.

A distinctive feature of our analysis compared with previous studies^11,30^ examining the predictive utility of GRS for CHD is that the best predictive model was achieved here with SNPs that did not necessarily reach genome-wide or even statistical significance in previous GWA studies. Genome-wide SNP models have been applied successfully to other heritable human traits which seem to follow an “infinitesimal” genetic architecture, such as height^20^. These results highlight the differing goals of GWAS and of genomic prediction: the stringent detection of causal genetic variants involved in the disease process versus the construction of a model that robustly and maximally predicts future disease. While stringent procedures for minimizing the false positive rate of associated loci in GWAS are appropriate, these concerns are less relevant in construction of GRSs, especially when there are a large number of weakly correlated SNPs^22^ and when rigorous internal and external validation is performed.

While population stratification is a potential confounder of genomic prediction studies, our use of a large worldwide multi-ethnic meta-analysis to develop the GRS together with two fully independent prospective validation datasets and three independent case/control datasets minimizes this potential. Our GRS was constructed from the CARDIoGRAMplusC4D stage-2 meta-analysis and the FINRISK and FHS individuals are both independent of that study and of broadly European ancestry; thus it is unlikely that the GRS is substantially confounded by fine-scale population structure within these cohorts. Further, the LD-thinning threshold to maximize prediction was determined in the WTCCC and MIGen datasets prior to applying the GRS to ARGOS, FINRISK, or FHS. Nevertheless, for some measures, GRS gains were less pronounced in FHS than in FINRISK. This may partly be due to the different definitions of CHD in these studies or to differences in environmental exposures. In addition, the FRS was developed in the FHS, leading to potential over-estimation of its association with CHD in the current analysis. Hence, there may be benefit from future development of population-specific GRSs, which may yield greater predictive power within each population.

The association of the GRS with incident CHD was not substantially attenuated by traditional risk factors or clinical risk scores derived from these risk factors. Furthermore, the GRS was strongly associated with CHD in a study consisting purely of individuals with familial hypercholesterolemia. These results suggest that genomic risk exerts its effect on CHD risk through molecular pathways that are largely independent of the effects of cholesterol, systolic blood pressure, and smoking. A hitherto unresolved question has been the extent to which a family history would capture any information that may be provided through genetic analysis. Here, we clearly demonstrate the superior performance of direct genetic information over self-reported family history of CHD, which is often incomplete and imprecise in practice and is influenced by family size and competing causes of death.

While we observed improvements in discrimination (C-index) resulting from adding the GRS to the clinical risk scores when considering adults of all ages, the improvements were substantially higher in older individuals (>60 years old). Rather than being driven by age-related differences in the effect of the GRS, these results are likely driven by differences in the clinical risk scores between the younger and older adults. Unlike the GRSs, the clinical risk scores showed substantial differences across ages, driven by temporal changes in the underlying risk factors as well as age itself. Beyond the aims of identifying older adults with high CHD risk, the invariance of genomic risk makes it particularly useful for CHD risk prediction earlier in life, in young adulthood or before, when traditional risk factors are typically not measured and less likely to be informative of risk later in life.

Stratifying individual baseline smoking, systolic blood pressure, and total cholesterol levels measures into genomic risk groups revealed substantial differences in cumulative risk patterns. Importantly, this demonstrates that improved lifestyle may compensate for the innate increased CHD risk captured by the GRS. For men with high genomic risk, modifiable risk factors showed large effects on cumulative CHD risk. For women, the observed impacts of smoking, systolic blood pressure, and total cholesterol were low or not detectable in the low genomic risk group, particularly in FINRISK, however, we could not determine whether this was due to inadequate statistical power or other biological effects and further studies in larger cohorts of women are necessary to determine any clinical implications.

Given recent advances and reduced cost of genotyping microarrays and sequencing-based technologies and their cost efficiency, determination of genome-wide SNP variants (including the 49,310 SNPs used here) is no longer beyond the realm of clinical application. In this context, our results, if validated in further studies and across different populations, suggest a potential paradigmatic shift in the current CHD screening strategy which has existed for over 40 years - namely determination of genomic risk at an early stage with screening later in life through traditional clinical risk scores to complement background genomic risk. Based on early genomic risk stratification, individuals at higher risk may benefit from earlier engagement with nutritionists, exercise regimes, smoking cessation programs or be initiated early on medical interventions such as statin therapy or blood-pressure lowering medications to minimize future CHD risk. In this context it is notable that Mega et al. ^30^ recently demonstrated that the GRS of 27 CHD-associated SNPs better predicted which individuals would benefit most, both in relative and absolute terms, from statin treatment. In a study of type 2 diabetes, Florez et al. ^31^ has shown that the effects of increased genetic susceptibility to disease can be ameliorated by lifestyle (diet and exercise) and therapeutic (metformin) interventions. Similar possibilities exist for CHD, whereby early targeted prevention strategies based on genomic CHD risk may be implemented well in advance of clinical risk scores attaining predictive capacity at later ages^32^. Such early risk stratification will offer increased efficiency in allocating both therapeutic resources and lifestyle modifications with the potential for subsequent delay of onset of traditional risk factors and incident CHD risk.

While our study demonstrates both the independent and incremental predictive power provided by our GRS, it is important to note that even when combined with such scores, the overall positive predictive value still remains modest for an acceptable negative predictive value (**Supplementary Figure S10**). Furthermore, despite overall improved reclassification of 10 year risk, some individuals who went on to develop an incident event were reclassified at a lower risk by the addition of the GRS compared to their initial classification using a clinical score (**Table 3** and **Supplementary Table S4**), emphasizing the limitations of the current GRS. In this context, it should be noted that our GRS based on a starting list of 79,128 common SNPs tested by the CARDIOGRAMplusC4D consortium could be further improved. Future studies that construct GRSs using increased sample sizes and capturing the full spectrum of common and rare variants^9,33^ will likely provide additional gains in prediction and risk stratification.

In summary, this study has demonstrated the potential clinical utility of genome-scale GRS for CHD, both for early identification of individuals at increased CHD risk and for complementing existing clinical risk scores.

## Acknowledgements

This study was supported by the National Health and Medical Research Council (NHMRC) of Australia (grant no. 1062227) and the National Heart Foundation of Australia. GA was supported by an NHMRC Peter Doherty Early Career Fellowship (no. 1090462). MI was supported by a Career Development Fellowship co-funded by the NHMRC and the National Heart Foundation of Australia (no. 1061435). VS was supported by the Finnish Foundation for Cardiovascular Research. NJS holds a Chair funded by the British Heart Foundation and is a UK National Institute for Health Research Senior Investigator. The Myocardial Infarction Genetics (MIGen) Consortium Study was funded by the National Heart, Lung, and Blood Institute of the United States National Institutes of Health (R01 HL087676). Genotyping was partially funded by The Broad Institute Center for Genotyping and Analysis, which was supported by grant U54 RR02027 from the National Center for Research Resources. This study makes use of data generated by the Wellcome Trust Case-Control Consortium. A full list of the investigators who contributed to the generation of the data is available from www.wtccc.org.uk. Funding for the project was provided by the Wellcome Trust under award 076113 and 085475. The research leading to these results has received funding from the European Union Seventh Framework Programme (FP7/2007-2013) under grant agreement no. 261433 (Biobank Standardisation and Harmonisation for Research Excellence in the European Union — BioSHaRE-EU). We are grateful to the CARDIoGRAMplusC4D consortium for making their large-scale genetic data available. A list of members of the consortium and the contributing studies is available at www.cardiogramplusc4d.org. We thank Julie Simpson, Melbourne School of Population Health and Global Health (University of Melbourne), for advice regarding survival analyses.

